# Characterization of Hippo Signaling Components in the Early Dorsal Pancreatic Bud

**DOI:** 10.1101/2024.10.26.619721

**Authors:** Neha Ahuja, Caitlin Maynard, Tyler Bierschenck, Ondine Cleaver

## Abstract

All pancreatic lineages originate from a transitory structure known as the multipotent progenitor epithelium (MPE), which is a placode formed via epithelial stratification. Cells within the MPE undergo *de novo* lumenogenesis to give rise to an epithelial plexus, which serves as a progenitor niche for subsequent development of endocrine, ductal and acinar cell types. Recent evidence suggests that Hippo signaling is required for pancreatic cell differentiation, but little is known about the function of Hippo signaling in the development of the MPE. Here, we characterize the expression of YAP1, TAZ, and the Hippo regulators LATS1/2 kinases and MERLIN in early murine pancreatic epithelium, during epithelial stratification, plexus development and emergence of endocrine cells. We find that YAP1 expression is relatively low in the pancreas bud during stratification, but increases by E11.5. Intriguingly, we find that TAZ, but not YAP1, is expressed in early endocrine cells. We further find that MERLIN and LATS1/2 kinases are robustly expressed during the period of rapid stratification and become markedly apical at nascent lumens. To gain a better understanding of how Hippo signaling and lumen formation are connected, we analyzed the expression of Hippo signaling components in an *in vitro* model of lumen formation and found that they are dynamically regulated during lumenogenesis. Together, our results point to a relationship between Hippo signaling and lumen formation during pancreatic development.

**HIGHLIGHTS:**

- YAP1 expression in the early mouse pancreatic anlagen is low until approximately E11.5, when it becomes localized to cell nuclei in multipotent progenitor cells. At E14.5, we find nuclear YAP1 in ductal cells.
- YAP1 is not expressed in early and midgestation endocrine cells. By contrast, TAZ is expressed in first transition endocrine cells.
- Hippo regulators MERLIN and LATS1/2 kinases are robustly expressed in the early pancreatic bud by E10.5. Both MERLIN and LATS1/2 exhibit strong apical localization in epithelial cells at nascent microlumens.
- Using *in vitro* models of *de novo* pancreas lumen formation, we show that YAP1 nuclear localization is high in early phases of lumen formation and gradually decreases as lumens matures.

## INTRODUCTION

Pancreatic development can be broadly classified into two stages: the primary transition and the secondary transition. During the primary transition, which begins at E8.75 in mouse embryos, the pancreatic placode emerges as thickening of the endodermal epithelium. During this period, the first endocrine cells also emerge from the epithelium.^1^ This placode then undergoes rapid proliferation and stratification to give rise to the E10.5 dorsal pancreatic bud, which consists of multipotent progenitor epithelial (MPE) cells from which all pancreatic cell lineages arise. Cells within the MPE undergo *de novo lumenogenesis*, which requires acquisition of cell polarity and cellular shape changes, to give rise to the lumenal network of the pancreas.^2^ At the onset of the secondary transition (approximately E12.5), the pancreatic epithelium consists of tip and trunk domains. The tip domain is highly proliferative and will give rise to the pancreatic acinar cells.^3,4^ The trunk domain consists of bipotent progenitors, which will give rise to the ductal and endocrine cells. As the lumenal network expands, the pancreatic epithelium concomitantly destratifies and returns to a simple monolayer epithelium by birth.

The MPE is a critical transitional tissue that develops due to the rapid stratification of endodermal cells from the gut tube. Experimental reduction in the pool of multipotent progenitors through E9.5-E11.5 has a dramatic effect on pancreatic size at E18.5, suggesting that proper allocation of MPE cells is critical for overall pancreatic development.^5^ Despite its importance, little is known about the cellular and molecular mechanisms that govern MPE development. Recent evidence has shown that Hippo signaling pathway is critically required for the development of a variety of organ systems,^6^ including the pancreas.^7^ In the canonical Hippo pathway, a kinase cascade is activated whereby the core kinases MST1/MST2 phosphorylate the LATS1/2 kinases. LATS1/2, in turn, inhibit the translocation of transcriptional regulators YAP1 and TAZ to the nucleus, abrogating transcription of their target genes. Additionally, Hippo signaling is regulated by MERLIN. MERLIN recruits LATS kinases to the cell membrane where they are then activated by MST1/2.^8^ Together, these molecules form a signaling cascade that regulates organ size and cell proliferation.^9^

There is significant interest in understanding the role of the Hippo signaling pathway in pancreatic development. At E10.5, YAP1 is present in pancreatic epithelium, and nuclear YAP1 is progressively enriched in productal epithelium.^10^ Acinar cells display predominant cytoplasmic (inhibited) YAP1, while islet endocrine cells are completely devoid of any YAP1 expression. This pattern of nuclear YAP1 in ductal cells and cytoplasmic YAP1 in acinar cells persists through adulthood.^10^ Indeed, YAP1 is a critical negative regulator of endocrine cell differentiation in both *in vivo* and *in vitro* models.^11,12^ We previously showed that LATS1 and LATS2 kinases are localized to the apico-junctional membrane of pancreatic epithelial tubules and that pancreatic knockout of both *Lats1* and *Lats*2 kinases (*LATS1/2*^*PancDKO*^) prevents proper differentiation of all pancreatic cell lineages.^7^ This phenotype was accompanied by dysregulation of apical-basal polarity proteins and cell adhesion molecules, and by partial-EMT (ectopic expression of mesenchymal markers in epithelium).^7^

Despite the critical importance of the Hippo pathway in regulating pancreatic morphogenesis and cellular differentiation, little is known about the requirement for Hippo pathway components during the initial phases of epithelial stratification and MPE development. In this study, we set out to characterize the expression of Hippo pathway components in the early pancreatic epithelium. We first see that YAP1 localization is predominantly non-nuclear during epithelial stratification, but that nuclear YAP1 increases by approximately E11.5 in the mouse embryos. We further find that YAP1 is not present in either primary or secondary transition endocrine cells. Intriguingly, we see that TAZ is present in the primary transition endocrine cells, suggesting that distinct mechanisms exist to regulate expression of these two critical transcriptional regulators. In addition, we evaluate expression and localization of key Hippo regulators, MERLIN and LATS1/2 kinases in the embryonic epithelium, since both molecules are able to directly interact with YAP1.^13^ We find that both MERLIN and LATS1/2 kinases are present in the pancreatic bud by E10.5, and both show robust apical localization at nascent lumens. Given striking localization of these Hippo pathway components to the apical membrane of newly forming lumens, we investigated the localization of Hippo pathway components during lumenogenesis using a 3-D sphere forming assay. We find that YAP1 is initially robustly nuclear during the initial phases of lumen formation and becomes more cytoplasmic as the lumen matures. Together, our results provide insight into the expression pattern of Hippo pathway components during early pancreatic development and point to an intrinsic relationship between lumen formation and Hippo signaling.

## MATERIALS AND METHODS

### Mice and embryo handling

Experiments were performed in accordance with protocols approved by the UT Southwestern Medical Center IACUC. E8.5 – E16.5 embryos were dissected and fixed in 4% PFA/PBS for overnight hours at 4°C.

### Tissue embedding and Immunofluorescence

Fixed pancreatic issue were washed in PBS and embedded in paraffin as previously described.^2,7^ In brief, tissue was gradually dehydrated into 100% molecular biology grade ethanol and then washed in xylene twice for 10 minutes per wash. Tissue was then placed in paraffin for at least 3 1-hour long washes, and then embedded in a paraffin cassette. Tissue was then sectioned at 10 µm on a microtome.

Sections were then rehydrated through an ethanol gradient, and were washed in 3x in PBS for at least 5 minutes per wash. Sections were permeabilized for 12 minutes 0.3% Triton in PBS. We performed antigen retrieval with either Buffer A (Electron Microscopy Sciences, cat number 50-311-77) for nuclear antigens or Buffer B (Electron Microscopy Sciences, cat number 50-311-78) for cytoplasmic antigens using a pressure cooker. Slides were then blocked for 1 hour in Cas Block (Thermo Fisher, cat number 008120). Primary antibody incubations were done at 4°C overnight (for catalog numbers and dilutions, see Supplementary Table S1). Slides were then washed in PBS, incubated in secondary antibody for an hour at room temperature. Slides were then washed in PBS and mounted using Flouromount Mounting media with DAPI. Images were obtained using a Nikon A1R confocal.

### Pancreatospheres

Pancreatospheres were made as previously described.^14,15^ In brief, CD1 wildtype E12.5 pancreata were dissected from embryos. The mesenchyme was removed by dispase treatment and manual dissection. Pancreata were then pooled into one Eppendorf tube and suspended in 0.1 mL of TrypLE (cat number 12604021). Pancreata were then incubated at 37°C for 5 minutes, and manually disaggregated by pipetting. TrypLE was neutralized by addition of 500µL of DMEM-F12/10% FBS (Thermo, cat number 12634010). Cells were spun down at 500g for 10 minutes at 4°C, diluted to 80 cells/uL in PEM Meda (DMEM-F12/10% FBS, 10µM rock inhibitor (Millipore Sigma, Y0503-1MG) 10ng/mL Fgf2 (Thermo Fischer, 450-33-100UG) and then diluted 1:3 in Matrigel. The cell suspension/Matrigel mixture was then pipetted onto 24-well coverglass-bottom plates (Mattek, cat number P24G-1.5-10-F) using wide bore tips and incubated at 37°C, 5% CO_2_ for 10 minutes to enable the Matrigel to cure. 500µL of PEM media was added to each well, and cells were incubated at 37°C, 5% CO_2_ until the appropriate time point. Spheres were fixed in 4% PFA in PBS at room temperature for 10 minutes. For IF, spheres were then washed 3 times in PBS for 5 minutes per wash, and then permeabilized in 0.1% NP-40 for 10 minutes. Spheres were then blocked in Cas block for 1 hour, and primary antibody incubations were performed overnight at 4°C. Secondary antibody incubations were performed at room temperature for between 1-2 hours, alongside 1:10000 DAPI incubation. Sphere were imaged on a Nikon Spinning Disk SORA at UT Southwestern Medical Center’s quantitative light microscopy core.

### Data Quantification and Statistical Analysis

All images shown are representative images from 2-3 biological replicates. For quantification of nuclear YAP1 localization on paraffin sections, the number of cells within the pancreatic epithelium with clear YAP1 expression in the DAPI+ nucleus was manually counted on each field of view. 2-3 sections were evaluated per embryo. For quantification of nuclear YAP1 intensity in spheres, nuclear fluorescence intensity was quantified by tracing the DAPI positive region of each cell and measuring the mean gray value of YAP1 fluorescence intensity using ImageJ Fiji (rep). Statistics were performed in Graphpad Prism 10, through either ANOVA or Student’s t-test as appropriate.

## RESULTS

### YAP1 expression is low during pancreatic stratification, but high by E11.5

To determine the spatiotemporal localization of YAP1 during the initial phases of pancreatic development (**Fig. 1A**), we performed immunofluorescence (IF) against YAP1 between E8.75 and E11.5. At E8.75, we observed weak YAP1 in the Pdx1+ pancreatic placode, but strong nuclear YAP1 in the adjacent gut tube (**Fig. 1B-C**). By E10.5-E11.0, we detected nuclear YAP1 in a subset of pancreatic epithelial cells in the dorsal pancreatic bud **(Fig. 1D)**. At E11.5, we observed YAP1 immunoreactivity throughout the pancreatic epithelium. Some pancreatic epithelial cells displayed nuclear YAP1, while others showed cytoplasmic or no YAP1 expression **(Fig. 1E)**. Together, these results show that YAP1 is relatively low during pancreatic epithelial stratification, but increases expression towards the end of the primary transition.

**Figure 1.**
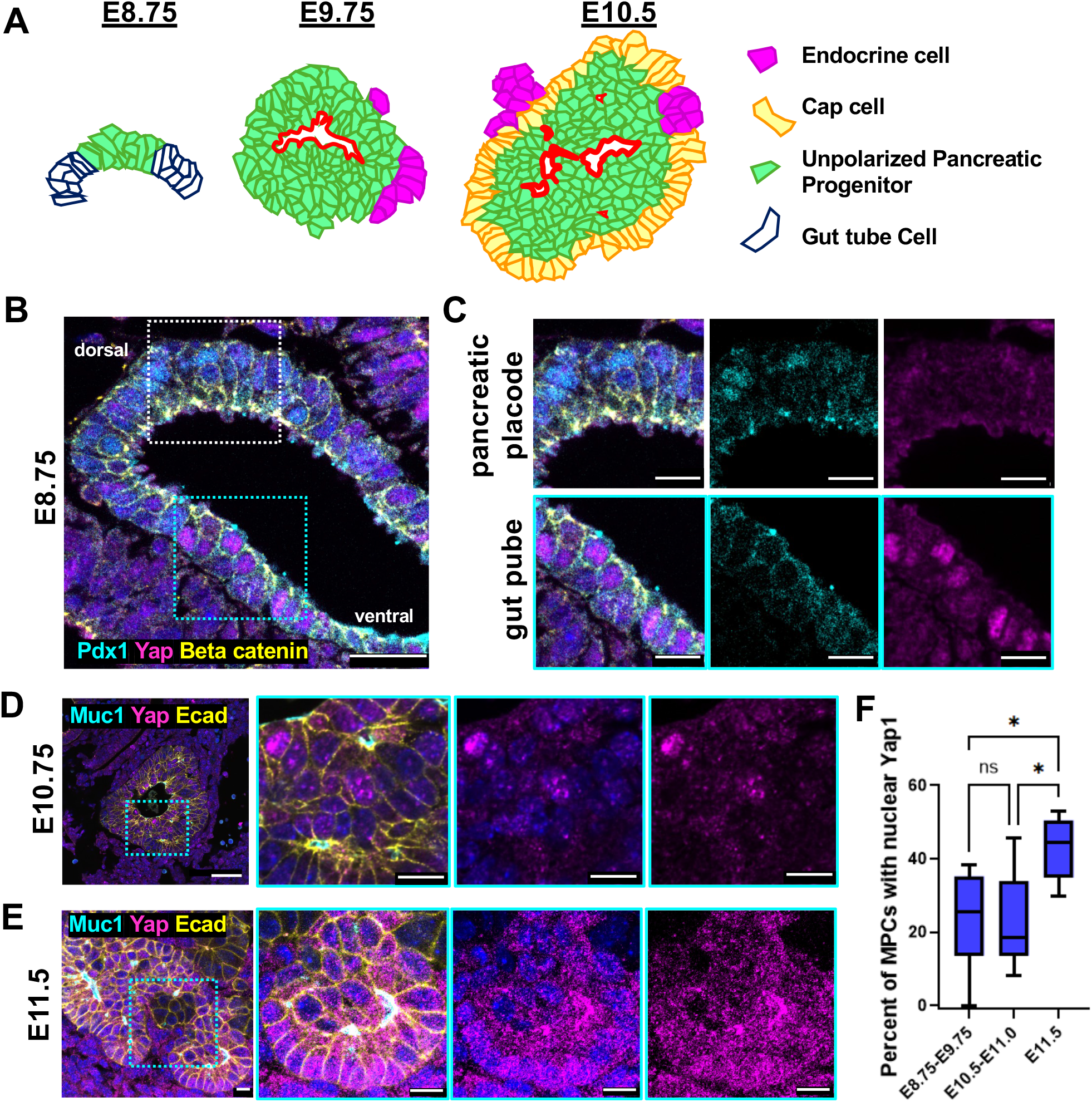
Pancreatic YAP1 expression becomes robust during the primary transition. **A)** Diagram of early pancreatic development. At 8.75, the pancreatic bud begins to form as an outgrowth of the gut tube. The dorsal pancreatic bud begins to undergo a period of rapid stratification. The first primary transition endocrine cells (primarily glucagon+ cells) emerge at ∼E9.75. At E10.5, the dorsal pancreatic bud consists of three cell types: 1) the outer columnar cap cells, the apolar inner body cells, and primary transition endocrine cells. Microlumen formation is robust by E10.75. **B, C)** YAP1 is not robustly detected in the E8.75 pancreatic primordia. **B)** E8.75 gut tube and pancreatic primordia in the embryo that is in the process of turning (dorsal and ventral indicated). White squares indicate regions shown in higher magnification in C. **C)** Cells that display nuclear Pdx1 (cyan) do not detectibly show YAP1 immunoreactivity (magenta). By contrast, intestinal epithelium display high levels of YAP1 nuclear immunoreactivity. **D)** At E10.75, select cells in the pancreatic epithelium begin to display YAP1 immunoreactivity (white arrow). **E)** By E11.5, YAP1 expression is robust. The majority of cells display nuclear YAP1, in both the cap and body compartments of dorsal pancreatic bud. **F)** Quantification of percent of MPCs that display nuclear YAP1 through time. N = 3 embryos analyzed between E8.75-E9.75, 4 embryos between E10.0 to E11.0, 3 embryos at E11.5. * indicated p <0.05, by Anova followed by Dunnet’s test for multiple comparison. Scale bar indicates 10 µm.

### YAP1 is not present in endocrine cells, but TAZ is present in primary transition endocrine cells

To investigate if YAP1 displayed cell-type specific expression in the early pancreas, we performed IF with endocrine costains at E10.5 and E14.5. At E10.5, we did not detect YAP1 expression in Glucagon-positive early first transition endocrine cells **(Fig. 2A, red arrows)**, though we were able to detect some expression in the pancreatic epithelium **(Fig. 2A, arrowhead)**. Consistent with previous reports,^11^ we found that YAP1 expression was absent from the secondary transition endocrine cells at E14.5 **(Fig. 2B, red arrow)**. To verify the lack of Hippo-dependent transcriptional activity in these early endocrine cells, we evaluated Hippo-effector TAZ. Intriguingly, we found that TAZ was present in early primary transition endocrine cells **(Fig. 2C, arrows)** but was lower in secondary transition endocrine cells **(Fig. 2D, red arrows)**. We saw robust expression of TAZ within the ductal epithelium at E14.5 (**Fig. 2D, white arrowhead)**. These results suggest that YAP1 and TAZ are differentially expressed in primary and secondary transition endocrine cells.

**Figure 2.**
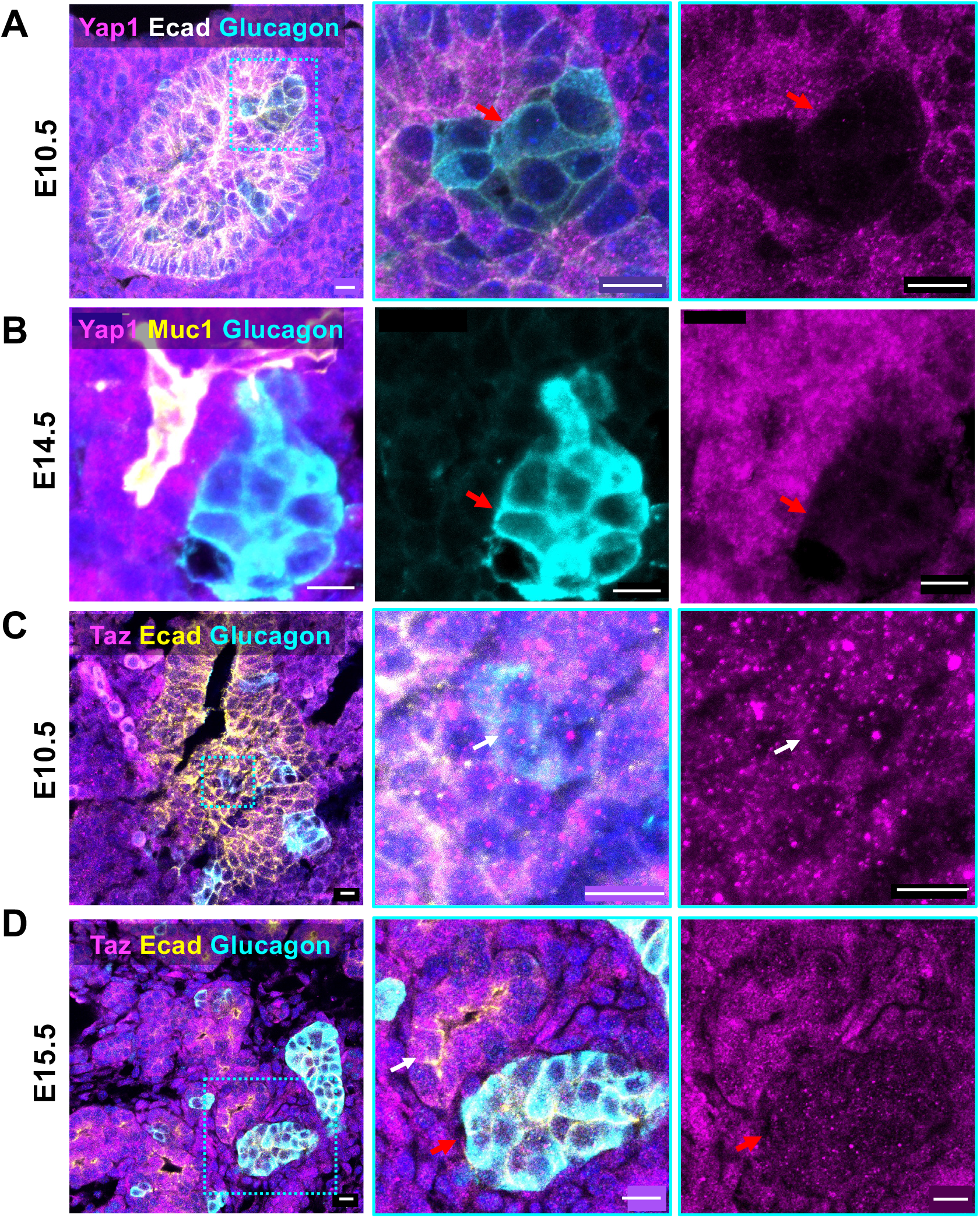
TAZ, but not YAP1, is expressed in primary transition endocrine cells. **A)** YAP1 is not expressed in primary transition endocrine cells. IF of E10.5 pancreatic bud. Note YAP1 immunoreactivity (magenta) is absent in the glucagon positive cells (red arrow). Representative image from n=3 pancreata. **B)** YAP1 is not expressed by secondary transition endocrine cells at E14.5. Representative image from n=3 pancreata. Red arrow indicates glucagon positive cells. **C)** TAZ is expressed in primary transition endocrine cells. IF of E10.5 pancreatic bud. (white arrow). Representative image from n=2 pancreata. **D)** TAZ is expressed by secondary transition endocrine cells (red arrow), though not as strongly as the ductal epithelial cells (white arrow) at E14.5. Representative image from n=3 pancreata. Scale bar indicates 10 µm.

### MERLIN and pLATS1/2 are apically localized in the pancreatic epithelium during the earliest stages of pancreatic development

To investigate how dynamic expression of Yap and TAZ may be achieved, we evaluated key Hippo upstream regulators LATS1/2 kinases and MERLIN. Lats/12 kinases directly phosphorylate YAP1/TAZ, sequestering them in the cytoplasm. MERLIN can interact with Hippo signaling in two ways: first, it can promote the phosphorylation of LATS1/2 kinases, thereby activating them; and second, MERLIN can directly bind to YAP1 and promote its export from the nucleus.

We first investigated the presence of phosphorylated LATS1/2 kinases (pLATS1/2) and MERLIN in the early pancreatic epithelium through IF using an antibody that recognizes the phosphorylated (active) form of both LATS1/2 kinases. As we previously showed, pLATS1/2 kinases are present in the early pancreatic bud with strong immunoreactivity by E9.75 in foci within the stratified epithelium **(Fig. 3A)**. ^7^ We investigated the subcellular localization through high resolution confocal microscopy, and found pLATS1/2 was primarily localized the apical junctional complex by E9.75 (**Supp Fig. 1A)**. Similarly, we found that MERLIN was present in the early pancreatic bud **(Fig. 3B)**. In cells that formed a lumen, we found that MERLIN was strongly apically localized **(Fig. 3B**, white arrows). In regions that were still stratified, MERLIN was expressed at the epithelial cell-cell interface, as indicated by colocalization with E-cadherin **(Fig. 3B**, blue arrow). By E11.5, we found that MERLIN was localized near mucin1, indicating that MERLIN is predominately localized the apical membrane, around nascent lumens (**Supp Fig. 1B**). To investigate if pLATS1/2 and MERLIN are expressed in primary transition endocrine cells, we evaluated localization of pLATS1/2 and MERLIN in glucagon (gluc) positive cells at E11.5. We found that pLATS1/2 displayed punctate expression in early endocrine cells, and that levels were lower than those at lumens **(Supp Fig. 1C)**. MERLIN was expressed along the cell periphery in these early endocrine cells **(Supp Fig. 1D)**. Together, these data indicate that Hippo regulators MERLIN and LATS1/2 kinases are expressed in early pancreatic epithelium in both multipotent progenitors and primary transition endocrine cells.

**Figure 3.**
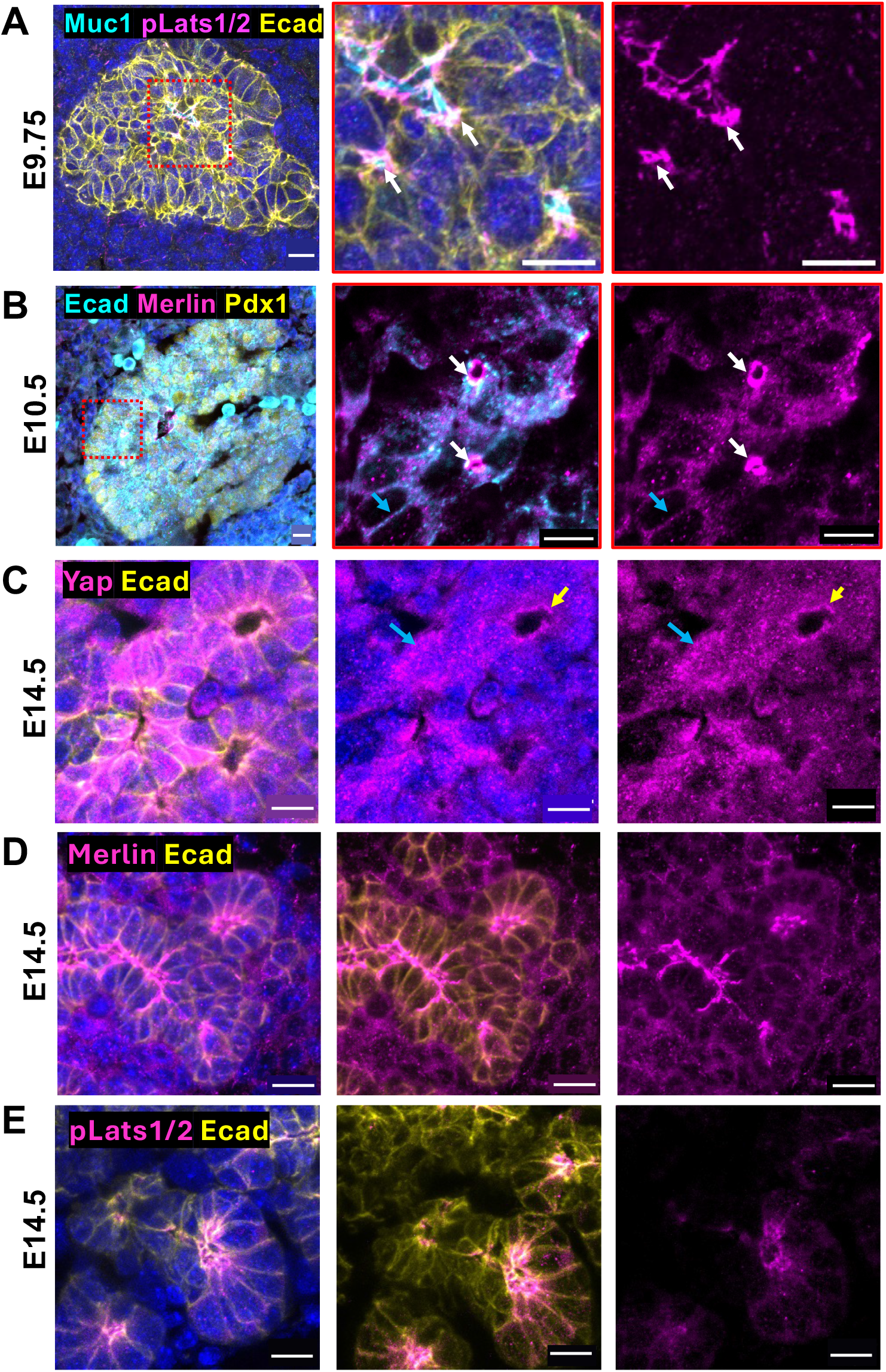
LATS1/2 kinases and MERLIN – critical regulators of Hippo signaling – are expressed both in the primary transition and secondary transition of pancreatic development. **A)** pLATS1/2 immunoreactivity is present by E9.75 in the early pancreatic bud. pLATS1/2 is present at nascent lumens, indicated by mucin1 staining (white arrows). MERLIN immunoreactivity is present by E10.5 in the early pancreatic bud. MERLIN is present at E-cadherin junctions (cyan arrow) and becomes enriched at the apical surface (white arrows). **C)** nuclear YAP1 is present in the E14.5 ductal epithelium (cyan arrow). In the acinar cells, YAP1 appears to cytoplasmic (yellow arrow). **D)** MERLIN and **E)** pLATS1/2 are present throughout the pancreatic epithelium at E14.5, primarily apically localized. For **A-B)** Images shown are representative through 2 embryos. For **C-E)** Images shown are representative of at least 3 embryos. Scale bar indicates 10 µm.

To determine if Hippo pathway components are expressed in more mature cell types, we investigated the localization of YAP1, pLATS1/2 and MERLIN in the E14.5 embryonic epithelium. Consistent with previous reports,^10^ we found that YAP1 was nuclear in ductal epithelium (**Fig. 3C, blue arrow**). In acinar epithelium, YAP1 did not appear to be enriched in the DAPI positive nucleus, and showed apical enrichment **(Fig. 3C, yellow arrow)**. We find that MERLIN and pLATS1/2 are present in both the ductal **(Fig. 3D)** and acinar epithelium **(Fig. 3E)**. Intriguingly, both MERLIN and pLATS1/2 are apically localized at this stage. Together these results show that pLATS1/2 and MERLIN are present in both ductal and acinar pancreatic epithelium, while YAP1 retains nuclear localization only in the ductal epithelium.

### YAP1 localization is dynamically regulated during *de novo* lumenogenesis

At E11.5, the stratified pancreatic epithelium is undergoing *de novo* lumenogenesis and interconnection between microlumens eventually gives rise to the epithelial plexus.^2^ The process of *de novo* lumenogenesis requires cells to undergo coordinated apical constriction, mediated by actomyosin contractility.^16^ Because the actomyosin machinery is implicated in the regulation of YAP1,^17^ we sought to understand if YAP1 is dynamically regulated during lumenogenesis by evaluating localization of YAP1 by IF at individual lumens within the E11.5 pancreatic epithelium.

We found that YAP1 localization was highly variable at an individual microlumens (**Fig. 4A, asterisk**) in the MPE, with select cells displaying robust nuclear YAP1 localization (**Fig. 4A, white arrows**) and cells displaying cytoplasmic/apical localization (**Fig. 4A, magenta arrows**). As a complementary approach to evaluating YAP1 nuclear vs cytoplasmic localization, we evaluated localization of inactive phosphorylated YAP1 through IF. Similar to YAP1, we found that there was significant variation in pYAP1 localization at any given microlumen (**Fig. 4B, asterisk**), with select cells displaying robust cytoplasmic pYAP1 immunoreactivity (**Fig. 4B, magenta arrows**) and other cells devoid of pYAP1 immunoreactivity (**Fig. 4B, white arrows**). Together, these data suggest that YAP1 localization and activity is dynamically regulated during lumen formation in the pancreatic epithelium.

**Figure 4.**
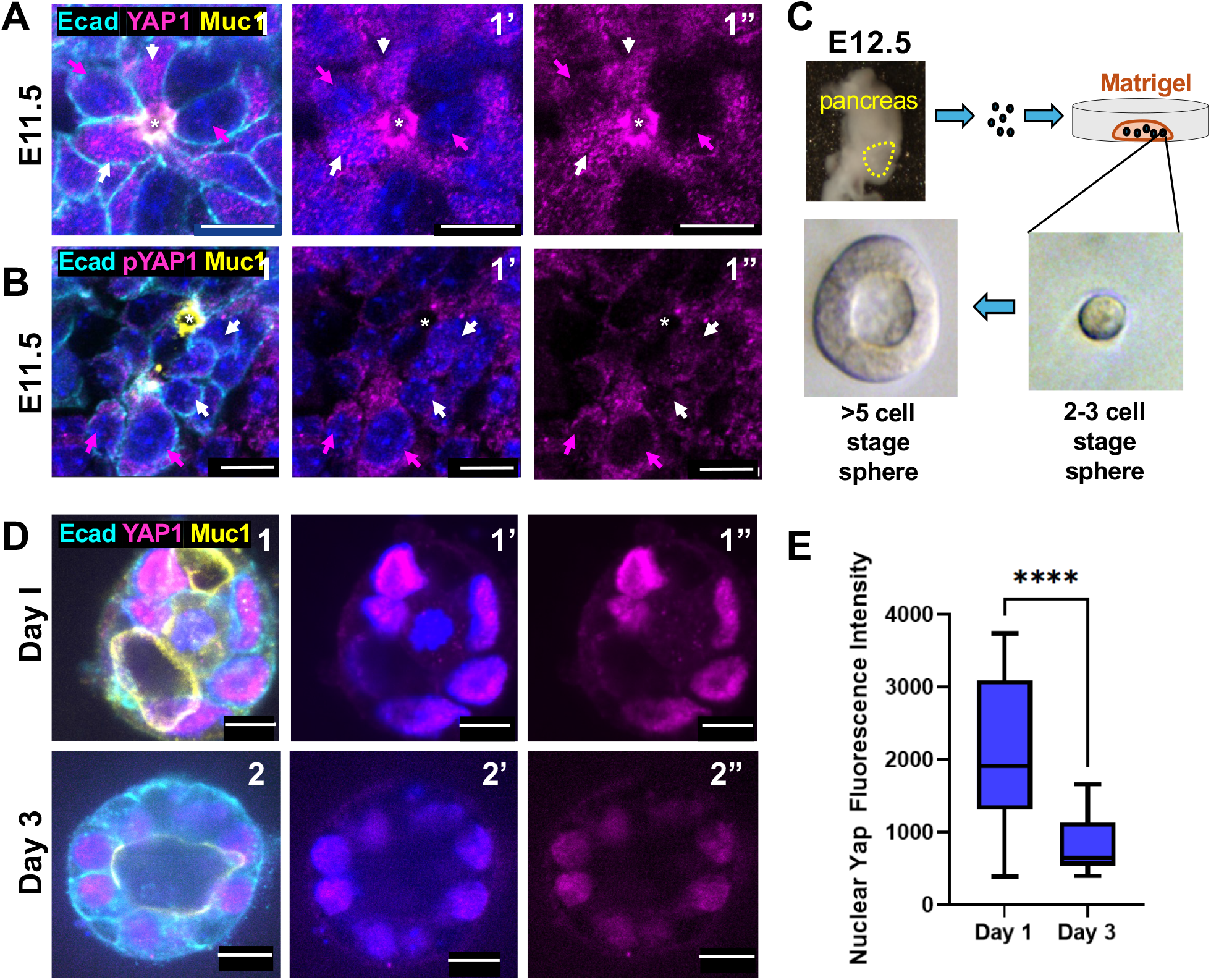
YAP1 is dynamically regulated during *de novo* lumen formation. **A)** YAP1 localization and expression is highly variable in the E11.5 pancreatic bud. High resolution imaging of a nascent lumen, indicated by mucin1 staining. Select cells participating in this lumen have robust nuclear YAP1 localization (white arrows), while some show distinctly cytoplasmic expression (magenta arrows). Lumen indicated by asterisk. **B)** IF of E11.5 YAP1, showing that some cells have inactivated YAP1. White arrows indicate cells that have low levels of pYAP1 immunoreactivity, magenta arrows indicate high levels of pYAP1 immunoreactivity. **C)** Schematic of sphere forming assay. Multipotent progenitor epithelial cells are isolated from the E12.5 pancreatic bud, dissociated into a single cell suspension, then cultured in Matrigel to promote sphere formation (as per ^15^). **D)** YAP1 is strongly nuclear at the onset of lumen formation (day 1), but becomes more cytoplasmic as lumen formation progress (day 3). **E)** Quantification of nuclear YAP1 intensity in pancreatosphere cells. N =3-5 spheres analyzed per time point, from 2 distinct replications. **** indicates p<0.0001, by student’s t-test. Scale bar indicates 10 µm.

To gain a better understanding of how YAP1 is regulated by lumen formation, we evaluated localization of YAP1 in an *in vitro* model of lumen formation, termed pancreatospheres.^15^ In this assay, MPEs from the E12.5 pancreatic epithelium are dissociated into single cells and then cultured in Matrigel. These cells then proliferate, acquire polarity, and form a lumen **(Fig. 4C)**, similar to pancreatic lumen formation *in vivo*.^*2*^ At day 1 of cell culture, cells are actively undergoing microlumen formation and often display multiple small luminal cavities **(Fig. 4D1)**. By day 3 of cell culture, cells form distinct lumens with clear apical polarity **(Fig. 4D2)**. This system enables us to track YAP1 activity from the initial stages of lumen formation until lumen maturation. At day 1 of culture, we observed strong nuclear YAP1 in these pancreatospheres **(Fig. 4D1-1”)**. There was a 2-fold decrease in nuclear Yap fluorescence intensity in spheres cultured for three days **(Fig. 4D2-2”)**, as compared to spheres with nascent lumens at 1 day of culture **(Fig. 4E)**. Together, these data suggest that YAP1 is transcriptionally active early during lumen formation and is sequestered in the cytoplasm as lumens mature.

## DISCUSSION

Early pancreatic development relies upon two primary morphogenetic paradigms: epithelial stratification and lumen formation.^18^ The precise regulation of these events is critical for both proper morphogenesis and endocrine differentiation; yet, the molecular cues that coordinate these initiating processes remain ill-defined. In this study, we characterize the expression of key Hippo signaling pathway components during two these two foundational phases of pancreas formation. Specifically, we show that YAP1 expression is relatively low during early pancreatic epithelial development, but becomes robust at the time when lumenogenesis is occurring. We further characterize the expression of YAP1 and TAZ in both primary and secondary transition endocrine cells, and intriguingly find that TAZ – but not YAP1 – is expressed in these early endocrine progenitors **(Fig. 5)**. This suggests that different mechanisms exist to regulate YAP1 and TAZ as the pancreas takes shape and also suggests the likelihood that YAP1 and TAZ have different roles. We show that key Hippo regulators MERLIN and LATS1/2 are present in both multipotent progenitors and early endocrine cells, underscoring a greater need for understanding how activity of these critical proteins are coordinated and regulated *in vivo*. Together, our observations are novel and suggest that Hippo signaling components are dynamically regulated during pancreatic development and cell fate acquisition.

**Fig 5:**
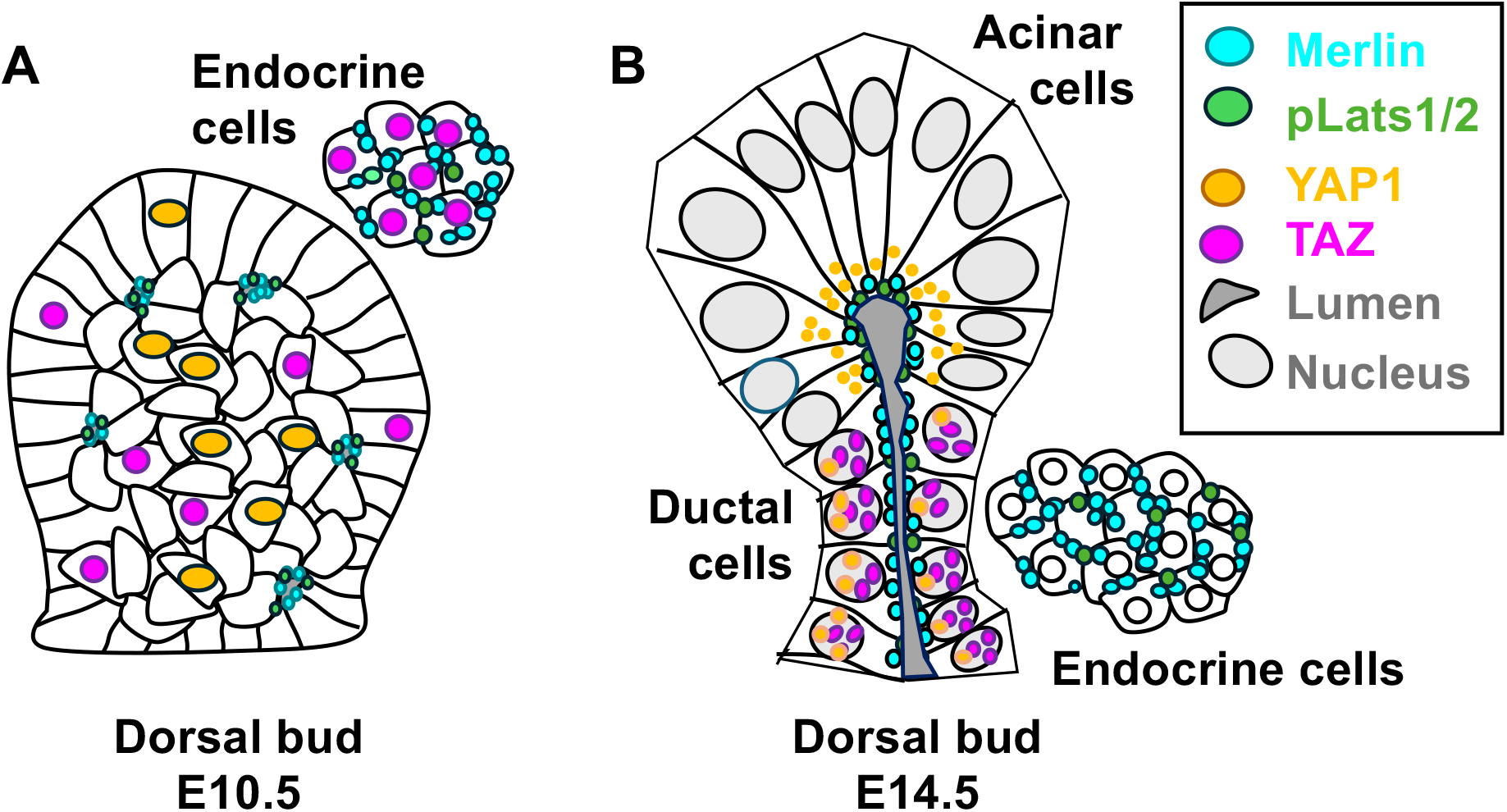
Summary of expression data. **A)** At E10.5, YAP1 and TAZ are expressed in a subset of cells within the MPE. TAZ, but not YAP1 is expressed in the early primary transition endocrine cells. LATS1/2 and MERLIN are present at the apical junctional and subapical regions, respectively. LATS1/2 and MERLIN are also present in endocrine cells. **B)** At E14.5, YAP1 is predominately nuclear in the ductal epithelium, and cytoplasmic in the acinar epithelium. Both YAP1 and TAZ have low to no expression in secondary transition endocrine cells. LATS1/2 and MERLIN are present in all pancreatic cell lineages.

The E10.5 pancreatic epithelium undergoes active cell proliferation, a process mediated by secreted signals from the surrounding mesenchyme.^19^ In other tissues, high levels of nuclear YAP1 are associated with high levels of epithelial cell proliferation; for example, overactivation of YAP1 in bronchial epithelial cells leads to lung hyperplasia.^20^ In our studies, however, we find that nuclear YAP1 levels are relatively low at E10.5, during the period of progenitor proliferation. Furthermore, overactivation of YAP1 through knockout of MST1/2 actually results in a smaller pancreas, indicating that overactivation of YAP1 is not sufficient to induce hyperplasia.^10^ Similarly, our previous studies show that inactivation of LATS1/2, which also results in overactivation of YAP1, results in a smaller embryonic pancreas.^7^ Does YAP1 signaling not play a role in regulating cell proliferation in pancreas as it does in other tissues? Or is the timing of cell proliferation and morphogenesis tightly controlled, leading to abrogation of further growth if dysregulated? These questions will require further studies. However, it is likely that other mechanisms exist to promote cell proliferation and epithelial stratification between E9.5-E10.5, perhaps downstream of paracrine signaling from the mesenchyme.

By contrast, we find that YAP1 nuclear levels increase dramatically by E11.5, a stage when the stratified pancreas epithelium is undergoing lumen formation. Lumen formation is a process that depends on cellular acquisition of polarity, and that has been linked to regulation of Hippo signaling and YAP1/TAZ subcellular localization.^21,22^ For example, in mouse pre-implantation embryos, cell division at the 8-cell blastula stage gives rise an apolar inner cell mass which are circumscribed by a ring of polar outer cells. The inner cell mass will give rise to the fetus and yolk sac, while other outer cells will give rise to the trophectoderm. Intriguingly, the subcellular localization of YAP1 is distinct in these two cellular compartments; YAP1 is strikingly nuclear in the polarized outer cells and cytoplasmic in the inner cell mass.^23^ This differential distribution of YAP1 is thought to mediated in part by apical polarity proteins. Knockdown of key polarity protein Pard6d reduced nuclear Yap in the outer cell mass by the 32-cell stage of mouse embryogenesis. This phenotype is due to disrupted localization of angiomotin (Amot), which is normally sequestered at the apical membrane by polarity proteins and directly interacts with LATS1/2 kinases to promote Hippo pathway activation.^24^ Work from *Drosophila* further supports the idea that Hippo signaling is regulated by apical polarization.^21^ Finally, and similarly, overexpression of Crumbs, an apical polarity determinant, results dramatic hyperplasia of the wing epithelium in a Hippo-dependent manner.^25^ Together, these many findings strongly suggest that apical polarity is a key determinant of Hippo activity.

In addition to polarity acquisition regulating YAP1 activity, it is also possible that a reciprocal relationship exists and that Hippo signaling regulates lumen formation. To date, the role of YAP1 in lumen formation has best been studied in the context of MDCK cysts, where knockdown of AmotL2 resulted in defects in lumen formation in a YAP1/TAZ dependent-manner.^26^ Instead of clear open lumens, cysts were filled with cells. This is reminiscent of our work with genetic deletion of LATS1/2 in the pancreas,^7^ which leads to early overgrowth of the epithelium and defects in lumen formation. Given that many organ systems that rely upon lumen formation -- including the cardiovascular system, pancreas, kidney and intestine – also require tight regulation of Hippo signaling,^7,27-29^ further investigation into the role of Hippo signaling during lumen formation is warranted.

Further studies have underscored that tight regulation of Hippo signaling is required for proper morphogenesis and cellular differentiation. Knockout of MST1/MST2 kinases from the pancreatic epithelium (*MST1/2*^*PancDKO*^) resulted in overactivation of YAP1 (as expected), and resulted in a loss of pancreatic cell mass. Though there was dysplasia of the pancreatic exocrine tissue, the endocrine cells were largely unaffected by loss of MST1/MST2.^10^ Intriguingly, *LATS1/2*^*PancDKO*^ display a drastically different phenotype – loss of LATS1/2 kinases from the pancreatic epithelium results in abrogates pancreatic morphogenesis and cellular differentiation. On a cellular level, *LATS1/2*^*PancDKO*^ displayed a partial epithelial-to-mesenchymal (EMT) transition, with robust induction of mesenchymal markers TAGLN and ACTA2 in mutant epithelium. Additionally, these mutant pancreata displayed defects in apical polarity, which was largely rescued by additionally knocking out *Yap1* from the *LATS1/2*^*PancDKO*^.^7^ Questions remain regarding why the phenotypes of *MST1/2*^*PancDKO*^ and *LATS1/2*^*PancDKO*^ are so markedly different, since they both result in significant YAP1 overactivation. One possibility is that LATS1/2 kinases have Hippo-independent roles to play in pancreatic organogenesis. In endothelial cells, LATS1/2 kinases directly phosphorylate AMOT, and facilitate cytoskeletal changes required for cell migration.^30^ It is also possible that LATS1/2 kinases may also direct cytoskeletal transitions required for lumenogenesis in a YAP1/TAZ-independent manner. Understanding the precise contribution of each Hippo signaling component to distinct steps in pancreas development, including possible roles in lumen formation, will likely yield novel insights into the molecular underpinnings of organogenesis across many different organ types.

Within the pancreatic epithelium, we –and others --find that Hippo signaling components are expressed in a cell-type specific manner. Specifically, we show that both YAP1 and TAZ are both present in multipotent progenitor populations in the E10.5 pancreas, and that YAP1 is not expressed in endocrine cells throughout development **(Fig 2)**. This raises an intriguing possibility that YAP1 activity may act as a cell fate determinant, and that high levels of YAP1 may prevent excess endocrinogenesis. Indeed, loss of *Yap1* from the early pancreatic epithelium resulted in an increase in insulin-producing β-cells and hypoglycemia by P4, suggesting that one function of YAP1 maybe to maintain the progenitor state.^11^ In this work, we show that TAZ is expressed in early primary transition endocrine cells, though its expression is decreased in secondary transition endocrine cells. More investigation needs to be done to understand the dynamics of TAZ in endocrine cell differentiation. Indeed, loss of *Taz* from the early pancreatic epithelium resulted in pancreata that displayed decreased islet size and β-cell mass, suggesting that TAZ is required for proper endocrine development.^31^ These differing phenotypes and expression patterns strongly suggest that YAP1 and TAZ have differential roles to play in pancreatic development. Future work should tease apart both what these individual proteins are doing and also identify mechanisms by which YAP1 and TAZ are differentially regulated.

Recent evidence has highlighted the role of mechanical stimuli in regulation of Hippo signaling (extensively reviewed in ^32-34^). In addition to cues from cell junctions and contact-inhibition, actomyosin contractility is thought to regulate the subcellular localization of YAP1.^17^ For example, in the wing imaginal disc epithelium, polarity protein aPKC tethers Hippo regulator Kibra at the junctional cortex.^25^ In conditions of low actomyosin tension, Kibra translocates away from cellular junctions towards the center of the cell, where it then promotes Yki (the *Drosophila* homolog of *Yap1/Taz*) inactivation.^35^ Additionally, actomyosin tension is thought to directly regulate MERLIN and trigger its translocation to the nucleus, where it works to inhibit YAP1 in epithelial cell culture.^13^ In murine kidneys, knockout of MERLIN from the ureteric bud lineage resulted in dilated, cystic epithelial tubules and defects in tip identity, showing that MERLIN is required for proper tube formation.^27^ We—and others—have shown that tight regulation of the actomyosin apparatus is critical for pancreatic morphogenesis and endocrine cell fate.^16,36,37^ Could actomyosin contractility and the Hippo pathway form a signaling axis that coordinates epithelial morphogenesis with cell fate? Future work will investigate how Hippo components localization and activity are modulated by actomyosin contractility and investigate impacts on downstream processes such as lumen formation and cell fate.

Altogether, this paper provides insight into the subcellular localization of key Hippo pathway molecules during early pancreas development. While the field has made much progress in understanding the role of this key pathway in development, many interesting questions, including broadly what the function of Hippo is signaling in early pancreatic development. We suspect that Hippo signaling is intrinsically tied to lumen formation, and indeed the process of lumen formation and the Hippo signaling pathway may in fact influence each other. Understanding the relationship between Hippo signaling and lumen formation is key to understand how pancreatic tubes are built, and will likely shed light on the molecular events that underly pancreatic morphogenesis and cellular differentiation.

## Abbreviations

IF: immunofluorescence
MPE: multipotent progenitor epithelium

## ACKNOWLEDGEMENTS

We would like to thank all members of the Cleaver lab for their constructive feedback. This work was supported by HL113498 and DK121408 to O.C, and 3-pdf-2023-1327-AN to N.A.

## Supplemental Information

**Supplemental Figure 1.**
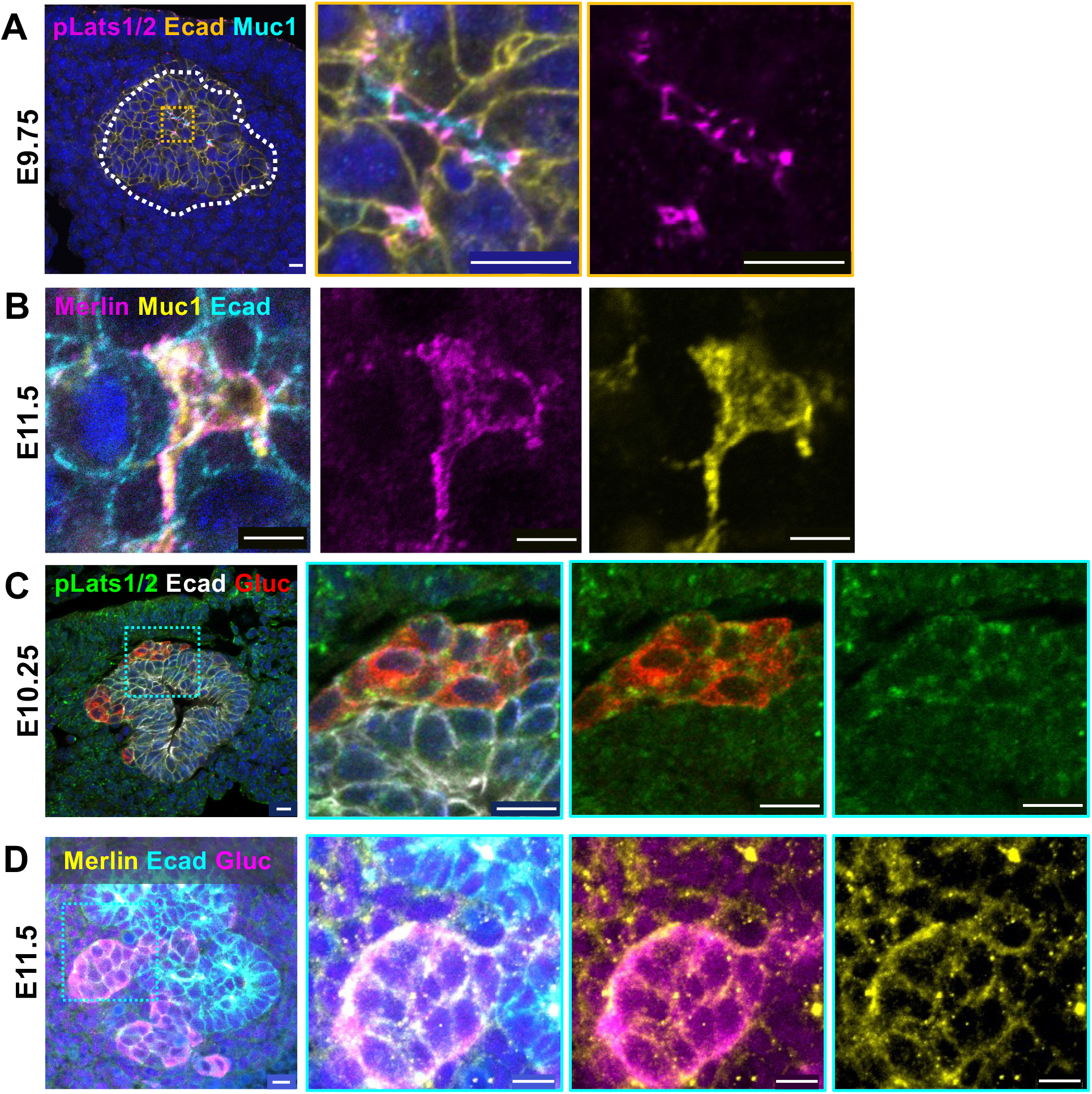
**A)** IF of pLATS1/2 in the early pancreatic epithelium. Insets show expression of pLATS1/2 are individual microlumens, and show pLATS1/2 restriction to the apical junctional region. **B)** MERLIN is localized to the apical domain at E11.5 pancreatic epithelium. **C)** IF of pLATS1/2 and Glucagon in the E10.25 pancreatic epithelium. pLATS1/2 is present in the early endocrine cells. **D)** IF of MERLIN and Glucagon in the E11.5 pancreatic epithelium. MERLIN is present in the early endocrine cells. Images are representative of n=2-4 embryos analyzed per stain. For A, C, and D, scale bar indicates 10 µm. For B, scale bar indicates 5 µm.

**Supplementary Table 1.**
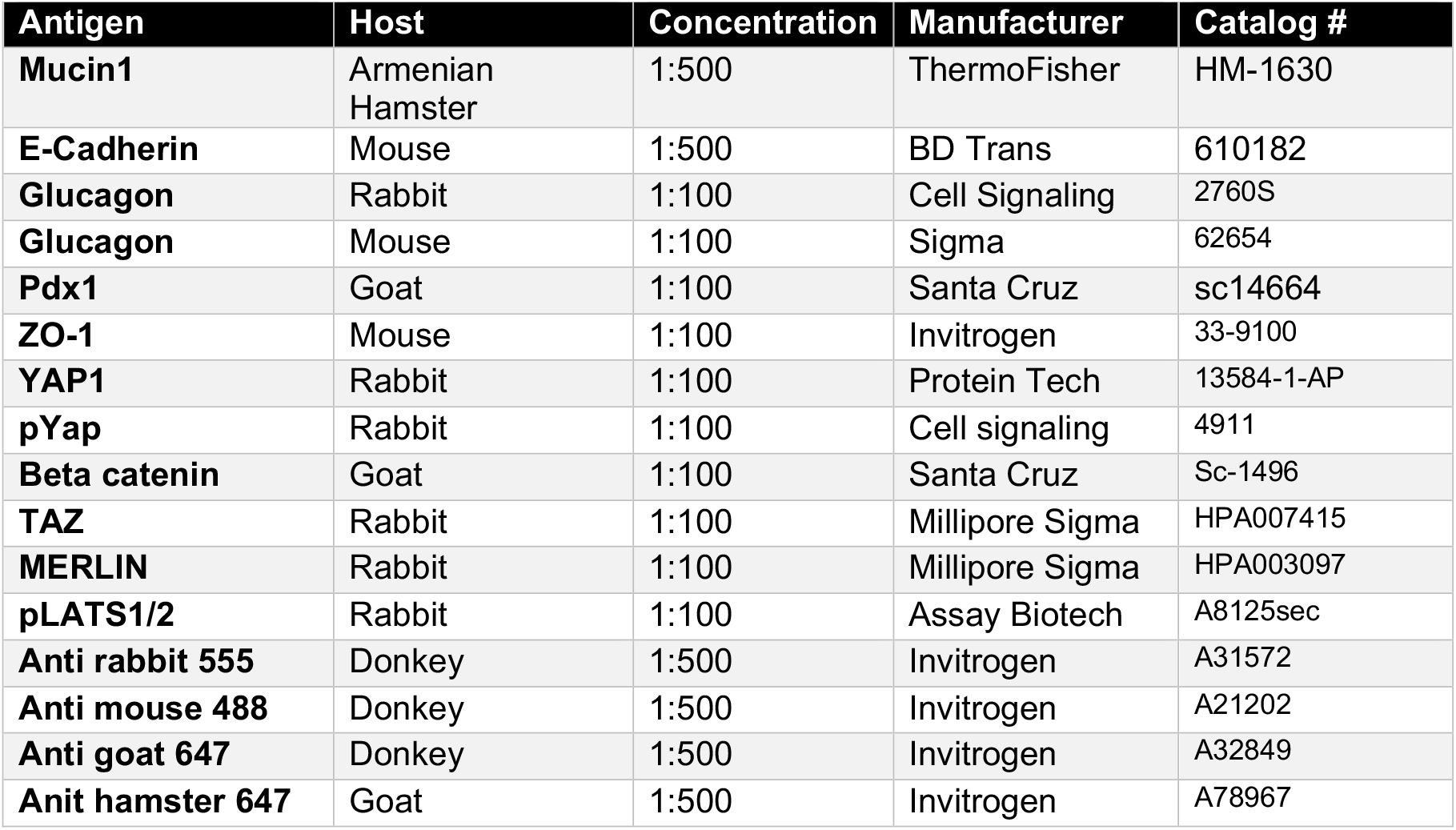
Antibodies used in presented studies.

| **Antigen** | **Host** | **Concentration** | **Manufacturer** | **Catalog #** |
| --- | --- | --- | --- | --- |
| **Mucin1** | Armenian Hamster | 1:500 | ThermoFisher | HM-1630 |
| **E-Cadherin** | Mouse | 1:500 | BD Trans | 610182 |
| **Glucagon** | Rabbit | 1:100 | Cell Signaling | 2760S |
| **Glucagon** | Mouse | 1:100 | Sigma | 62654 |
| **Pdx1** | Goat | 1:100 | Santa Cruz | sc14664 |
| **ZO-1** | Mouse | 1:100 | Invitrogen | 33-9100 |
| **YAP1** | Rabbit | 1:100 | Protein Tech | 13584-1-AP |
| **pYap** | Rabbit | 1:100 | Cell signaling | 4911 |
| **Beta catenin** | Goat | 1:100 | Santa Cruz | Sc-1496 |
| **TAZ** | Rabbit | 1:100 | Millipore Sigma | HPA007415 |
| **MERLIN** | Rabbit | 1:100 | Millipore Sigma | HPA003097 |
| **pLATS1/2** | Rabbit | 1:100 | Assay Biotech | A8125sec |
| **Anti rabbit 555** | Donkey | 1:500 | Invitrogen | A31572 |
| **Anti mouse 488** | Donkey | 1:500 | Invitrogen | A21202 |
| **Anti goat 647** | Donkey | 1:500 | Invitrogen | A32849 |
| **Anit hamster 647** | Goat | 1:500 | Invitrogen | A78967 |

## Notes

### Competing Interest Statement

The authors have declared no competing interest.

## REFERENCES

1. Benitez CM, Goodyer WR, Kim SK. Deconstructing pancreas developmental biology. Cold Spring Harb Perspect Biol. 2012;4. doi: 10.1101/cshperspect.a012401

2. Villasenor A, Chong DC, Henkemeyer M, Cleaver O. Epithelial dynamics of pancreatic branching morphogenesis. Development. 2010;137:4295–4305. doi: 10.1242/dev.052993

3. Kopp JL, Dubois CL, Hao E, Thorel F, Herrera PL, Sander M. Progenitor cell domains in the developing and adult pancreas. Cell Cycle. 2011;10:1921–1927. doi: 10.4161/cc.10.12.16010

4. Marty-Santos L, Cleaver O. Progenitor Epithelium: Sorting Out Pancreatic Lineages. J Histochem Cytochem. 2015;63:559–574. doi: 10.1369/0022155415586441

5. Stanger BZ, Tanaka AJ, Melton DA. Organ size is limited by the number of embryonic progenitor cells in the pancreas but not the liver. Nature. 2007;445:886–891. doi: 10.1038/nature05537

6. Zheng Y, Pan D. The Hippo Signaling Pathway in Development and Disease. Dev Cell. 2019;50:264–282. doi: 10.1016/j.devcel.2019.06.003

7. Braitsch CM, Azizoglu DB, Htike Y, Barlow HR, Schnell U, Chaney CP, Carroll TJ, Stanger BZ, Cleaver O. LATS1/2 suppress NFκB and aberrant EMT initiation to permit pancreatic progenitor differentiation. PLoS Biol. 2019;17:e3000382. doi: 10.1371/journal.pbio.3000382

8. Yin F, Yu J, Zheng Y, Chen Q, Zhang N, Pan D. Spatial organization of Hippo signaling at the plasma membrane mediated by the tumor suppressor Merlin/NF2. Cell. 2013;154:1342–1355. doi: 10.1016/j.cell.2013.08.025

9. Pan D. Hippo signaling in organ size control. Genes Dev. 2007;21:886–897. doi: 10.1101/gad.1536007

10. George NM, Day CE, Boerner BP, Johnson RL, Sarvetnick NE. Hippo signaling regulates pancreas development through inactivation of Yap. Mol Cell Biol. 2012;32:5116–5128. doi: 10.1128/MCB.01034-12

11. Mamidi A, Prawiro C, Seymour PA, de Lichtenberg KH, Jackson A, Serup P, Semb H. Mechanosignalling via integrins directs fate decisions of pancreatic progenitors. Nature. 2018;564:114–118. doi: 10.1038/s41586-018-0762-2

12. Rosado-Olivieri EA, Anderson K, Kenty JH, Melton DA. YAP inhibition enhances the differentiation of functional stem cell-derived insulin-producing beta cells. Nat Commun. 2019;10:1464. doi: 10.1038/s41467-019-09404-6

13. Furukawa KT, Yamashita K, Sakurai N, Ohno S. The Epithelial Circumferential Actin Belt Regulates YAP/TAZ through Nucleocytoplasmic Shuttling of Merlin. Cell Rep. 2017;20:1435–1447. doi: 10.1016/j.celrep.2017.07.032

14. Azizoglu DB, Braitsch C, Marciano DK, Cleaver O. Afadin and RhoA control pancreatic endocrine mass via lumen morphogenesis. Genes Dev. 2017;31:2376–2390. doi: 10.1101/gad.307637.117

15. Greggio C, De Franceschi F, Figueiredo-Larsen M, Gobaa S, Ranga A, Semb H, Lutolf M, Grapin-Botton A. Artificial three-dimensional niches deconstruct pancreas development in vitro. Development. 2013;140:4452–4462. doi: 10.1242/dev.096628

16. Marty-Santos L, Cleaver O. Pdx1 regulates pancreas tubulogenesis and E-cadherin expression. Development. 2016;143:1056. doi: 10.1242/dev.135806

17. Sun S, Irvine KD. Cellular Organization and Cytoskeletal Regulation of the Hippo Signaling Network. Trends Cell Biol. 2016;26:694–704. doi: 10.1016/j.tcb.2016.05.003

18. Jorgensen MC, Ahnfelt-Ronne J, Hald J, Madsen OD, Serup P, Hecksher-Sorensen J. An illustrated review of early pancreas development in the mouse. Endocr Rev. 2007;28:685–705. doi: 10.1210/er.2007-0016

19. Bhushan A, Itoh N, Kato S, Thiery JP, Czernichow P, Bellusci S, Scharfmann R. Fgf10 is essential for maintaining the proliferative capacity of epithelial progenitor cells during early pancreatic organogenesis. Development. 2001;128:5109–5117. doi: 10.1242/dev.128.24.5109

20. Lange AW, Sridharan A, Xu Y, Stripp BR, Perl AK, Whitsett JA. Hippo/Yap signaling controls epithelial progenitor cell proliferation and differentiation in the embryonic and adult lung. J Mol Cell Biol. 2015;7:35–47. doi: 10.1093/jmcb/mju046

21. Genevet A, Tapon N. The Hippo pathway and apico-basal cell polarity. Biochem J. 2011;436:213–224. doi: 10.1042/BJ20110217

22. Bryant DM, Mostov KE. From cells to organs: building polarized tissue. Nat Rev Mol Cell Biol. 2008;9:887–901. doi: 10.1038/nrm2523

23. Nishioka N, Inoue K, Adachi K, Kiyonari H, Ota M, Ralston A, Yabuta N, Hirahara S, Stephenson RO, Ogonuki N, et al. The Hippo signaling pathway components Lats and Yap pattern Tead4 activity to distinguish mouse trophectoderm from inner cell mass. Dev Cell. 2009;16:398–410. doi: 10.1016/j.devcel.2009.02.003

24. Hirate Y, Hirahara S, Inoue K, Suzuki A, Alarcon VB, Akimoto K, Hirai T, Hara T, Adachi M, Chida K, et al. Polarity-dependent distribution of angiomotin localizes Hippo signaling in preimplantation embryos. Curr Biol. 2013;23:1181–1194. doi: 10.1016/j.cub.2013.05.014

25. Chen CL, Gajewski KM, Hamaratoglu F, Bossuyt W, Sansores-Garcia L, Tao C, Halder G. The apical-basal cell polarity determinant Crumbs regulates Hippo signaling in Drosophila. Proc Natl Acad Sci U S A. 2010;107:15810–15815. doi: 10.1073/pnas.1004060107

26. Zhao B, Li L, Lu Q, Wang LH, Liu CY, Lei Q, Guan KL. Angiomotin is a novel Hippo pathway component that inhibits YAP oncoprotein. Genes Dev. 2011;25:51–63. doi: 10.1101/gad.2000111

27. Reginensi A, Enderle L, Gregorieff A, Johnson RL, Wrana JL, McNeill H. A critical role for NF2 and the Hippo pathway in branching morphogenesis. Nat Commun. 2016;7:12309. doi: 10.1038/ncomms12309

28. Marciano DK. A holey pursuit: lumen formation in the developing kidney. Pediatr Nephrol. 2017;32:7–20. doi: 10.1007/s00467-016-3326-4

29. Boopathy GTK, Hong W. Role of Hippo Pathway-YAP/TAZ Signaling in Angiogenesis. Front Cell Dev Biol. 2019;7:49. doi: 10.3389/fcell.2019.00049

30. Dai X, She P, Chi F, Feng Y, Liu H, Jin D, Zhao Y, Guo X, Jiang D, Guan KL, et al. Phosphorylation of angiomotin by Lats1/2 kinases inhibits F-actin binding, cell migration, and angiogenesis. J Biol Chem. 2013;288:34041–34051. doi: 10.1074/jbc.M113.518019

31. Jeong MG, Kim HK, Lee G, Won HY, Yoon DH, Hwang ES. TAZ promotes PDX1-mediated insulinogenesis. Cell Mol Life Sci. 2022;79:186. doi: 10.1007/s00018-022-04216-2

32. Chang YC, Wu JW, Wang CW, Jang AC. Hippo Signaling-Mediated Mechanotransduction in Cell Movement and Cancer Metastasis. Front Mol Biosci. 2019;6:157. doi: 10.3389/fmolb.2019.00157

33. Gumbiner BM, Kim NG. The Hippo-YAP signaling pathway and contact inhibition of growth. J Cell Sci. 2014;127:709–717. doi: 10.1242/jcs.140103

34. Ibar C, Kirichenko E, Keepers B, Enners E, Fleisch K, Irvine KD. Tension-dependent regulation of mammalian Hippo signaling through LIMD1. J Cell Sci. 2018;131. doi: 10.1242/jcs.214700

35. Tokamov SA, Nouri N, Rich A, Buiter S, Glotzer M, Fehon RG. Apical polarity and actomyosin dynamics control Kibra subcellular localization and function in Drosophila Hippo signaling. Dev Cell. 2023;58:1864–1879 e1864. doi: 10.1016/j.devcel.2023.08.029

36. Darrigrand J-F, Salowka A, Spagnoli FM. 2023. doi: 10.1101/2023.01.26.525717

37. Bankaitis ED, Bechard ME, Gu G, Magnuson MA, Wright CVE. ROCK-nmMyoII, Notch and Neurog3 gene-dosage link epithelial morphogenesis with cell fate in the pancreatic endocrine-progenitor niche. Development. 2018;145. doi: 10.1242/dev.162115

